# Aging interacts with tumor biology to produce major changes in the immune tumor microenvironment

**DOI:** 10.1101/2020.06.08.140764

**Authors:** Rossin Erbe, Zheyu Wang, Neeha Zaidi, Michael Topper, Stephen Baylin, Elizabeth M Jaffee, Hariharan Easwaran, Elana J Fertig

## Abstract

Advanced age is strongly correlated with both increased cancer incidence and general immune decline. The immune tumor microenvironment (ITME) has been established as an important prognostic of both therapeutic efficacy and overall patient survival. Thus, age-related immune decline is an important consideration for the treatment of a large subset of cancer patients. Current studies of aging-related immune alterations are predominantly performed on non-cancerous tissue, requiring additional study into the effects of age on tumor immune infiltration. We leverage large scale transcriptional data sets from The Cancer Genome Atlas and the Genotype-Tissue Expression project to distinguish normal age-related immune alterations from age-related changes in tumor immune infiltration. We demonstrate that while there is overlap between the normal immune aging phenotype and that of the ITME, there are several changes in immune cell abundance that are specific to the ITME, particularly in T cell, NK cell, and Macrophage populations. These results suggest that aged immune cells are more susceptible to tumor suppression of cytotoxic immune cell infiltration and activity than normal tissues, which creates an unfavorable ITME in older patients in excess of normal immune decline with age and may inform the application of existing and emerging immunotherapies for this large population of patients. We additionally identify that age-related increases in tumor mutational burden are associated with decreased DNA methylation and increased expression of the immune checkpoint genes *PDL1, CD80,* and *LAG3* which may have implications for therapeutic application of immune checkpoint blockade in older patients.

## Introduction

The association of cancer incidence with age is well established and the phenomenon of age-related immune decline has been recognized for even longer (Gardner, 1980). Mutations and epigenetic alterations are believed to accumulate with age and drive carcinogenesis (Tomasetti et al., 2017), (Horvath, 2013). Recent research has highlighted the specific changes that contribute to the general decline of the immune system that occurs as individuals age (Aw et al., 2007). Understanding the effect such alterations have on the anti-tumor immune response is critical for the informed development and application of immunotherapies to elderly patients.

Thus, systemic immune aging has received considerable attention in the context of its effect on cancer development and progression (Fulop et al., 2017). In particular, loss of T cell receptor (TCR) diversity (Britanova et al., 2014), decreased capacity of cytotoxic cells (Solana and Mariani, 2000), and increased inflammatory signaling (Franceschi et al., 2000) have been identified as age-related immune changes of potential relevance to cancer therapeutics and patient survival. Currently, these age-related changes in the immune system are identified predominantly in non-cancerous tissues. However, tumors actively alter immune cells and their immune microenvironment to promote disease progression and avoid targeting by the immune system, which has been identified as one of the major hallmarks of cancer (Hanahan and Weinberg, 2011). Therefore, additional aging-related shifts may occur in the immune tumor microenvironment (ITME) as a result of interactions between tumor immunosuppressive signaling and the immunosenescence phenotype.

In the last decade, the composition of the ITME has become a subject of intense study due to its association with overall survival and therapeutic efficacy, particularly that of immune checkpoint blockade (ICB) (Frankel et al., 2017). However, any shifts in the composition of the ITME itself that may occur with age have not yet been generally characterized, with the exception of some T cell populations in melanoma (Kugel et al., 2018). Merging the disparate research on cancer and aging can further distinguish whether age-related changes in non-cancerous tissues are recapitulated within the ITME. At present, ICB immunotherapy is less often used to treat elderly patients, due to concerns about efficacy and toxicity, despite the fact that the limited clinical trial data that exists suggests that they experience no reduced benefit as compared to younger patients (Kugel et al., 2018), (Elias et al., 2018), (Daste et al., 2017), (Jain et al., 2019). Agespecific characterization of the ITME is essential to understand these results and move forward with efforts to bring immunotherapy to this large group of cancer patients.

This study directly compares the impact of aging on immune response and infiltration in tumors to that of non-cancerous tissues using genomics data from large-scale consortium studies. We apply immune cell type deconvolution methods to bulk transcriptomics data from untreated tumors samples among patients of different ages from The Cancer Genome Atlas (TCGA) and to post-mortem non-cancer tissue samples from individuals of different ages from the GenotypeTissue Expression project (GTEx) in order to compare age-associated immune composition changes from within tumors and non-cancerous tissues. We identify decreases in overall T cell abundance in tumor samples that are not observed in systemic tissues, as well as an increase in macrophage abundance. Further, while NK abundance generally increases with age, this trend is not observed within the ITME. These cancer-specific changes are both poor prognostics based on TCGA overall survival data and the existing literature on the impact of these immune cell types in the ITME. These analyses suggest that not only do older cancer patients face the normal aspects of immune decline, but that there is a specific interaction between the senescent immune system and tumor signaling that produces a less favorable ITME for survival and therapeutic response. This represents an unappreciated interaction between tumor biology and the aging immune system that contributes to the worsening of clinical outcomes with increasing age and may suggest new treatment strategies for elderly patients. At the same time, we observe that increasing tumor mutational burden with age appears to lead to an epigenetically regulated increase in tumor expression of immune checkpoint receptors such as *PDL1, CD80,* and *LAG3,* which may make ICB an unexpectedly attractive therapeutic option for many older patients.

## Results

### Deconvolution of immune cell abundance in tumor samples from TCGA reveals an age-related decrease in T cell abundance and increase in macrophage abundance that is associated with decreased survival

The large number of primary tumor transcriptional profiles across disease subtypes available from TCGA provides a unique cohort to characterize the impact of age on the ITME. We apply the MIXTURE immune cell type deconvolution algorithm (Fernandez et al., 2019) to infer the absolute proportions of immune cell types from RNA-sequencing data derived from pan-cancer TCGA samples. We then fit a linear model with age, including cancer type (based on the 33 TCGA cancer-type projects included in the study) and patient sex as covariates, for each immune cell type (listed in Supplemental Table 1) to assess changes in immune cell infiltration as patients age. We find that overall T cell abundance significantly decreases with age in the ITME (q-value = 0.00175) (Figure 1A) while macrophages significantly increase in abundance (q = 4.45 x 10^-4^) (Figure 1B). Detectable changes in the infiltration of NK cells, Dendritic cells, B cells, and other myeloid populations do not occur with age pan-cancer (Figure 1C, Table 1).

**Figure 1–.**
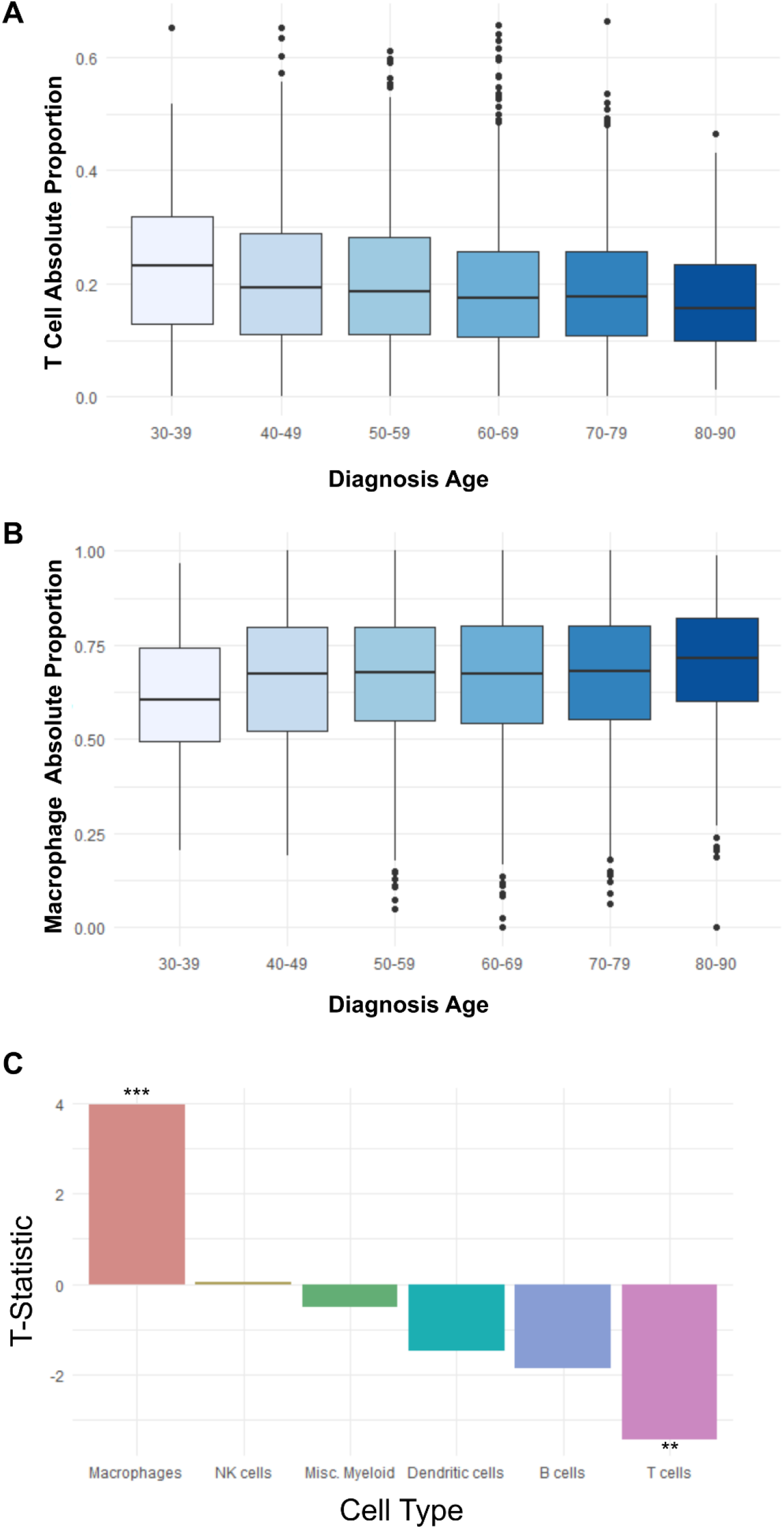
Macrophages increase and T cells decrease with age in the ITME. **A** Boxplots representing the pan-cancer change in T cell absolute proportion across different age groups from TCGA. A significant downward trend is observed based on a covariate-adjusted linear model **B** Boxplots representing the change in macrophage absolute proportion across age groups in TCGA. A significant upward trend is observed based on a covariate-adjusted linear model **C** Barplot of the t-statistics for the diagnosis age term of the linear models fit to each immune cell type from TCGA, including sex and cancer type as covariates. ** indicates an adjusted p-value less than 0.01 and *** indicates less than 0.001.

**Table 1.** Coefficients, statistics, p, and q-values for the diagnosis age term in the linear model fit for each immune cell type. Cancer type and sex were included as covariates for each of these models.

|  | **Estimate** | **Standard Error** | **t-value** | **p-value** | **q-value** |
| --- | --- | --- | --- | --- | --- |
| **T cells** | -0.0006 | 0.000175 | -3.44233 | 0.000585 | 0.001754 |
| **Macrophages** | 0.001075 | 0.000271 | 3.967909 | 7.42E-05 | 0.000445 |
| **B cells** | -0.00028 | 0.000149 | -1.87737 | 0.060566 | 0.121131 |
| **NK cells** | 3.82E-06 | 6.89E-05 | 0.055398 | 0.955826 | 0.955826 |
| **Dendritic cells** | -0.00015 | 9.77E-05 | -1.49017 | 0.136286 | 0.204429 |
| **Misc. Myeloid** | -4.97E-05 | 9.92E-05 | -0.50055 | 0.616722 | 0.740066 |

We additionally investigate age-related changes in the TCGA cohort at a finer cellular resolution of 22 immune cell subtypes from MIXTURE (listed in Supplemental Table 1) (Supplemental Figure 1). Naive B cells are found to significantly decrease in abundance with increasing age in the ITME (q-value = 0.0305). Although many known systemic immune changes occur with age in healthy tissues, this reduction in Naive B cells is the only statistically significant shift among immune cell subtypes in the ITME. Consistent with our previous analysis of the higher order cell types, the three macrophage subsets (M0, M1, and M2) are among the top four cell types that trend towards increasing abundance in age. Likewise, all T cell subsets (CD8, CD4, Follicular helper, and regulatory T cells) trend towards a decrease in abundance with age with the exception of gamma delta T cells.

To investigate the association of these age-related immune infiltration changes with patient survival, we fit a Cox proportional hazards model between overall survival and all immune cell types (both subtypes and higher order cell types), adjusting for patient age, sex, cancer type, and years of smoking (Supplemental Figure 2, Table 2). We observe that higher risk is significantly associated with lower T cell abundance (q = 5.38 x 10^-4^) and NK cell abundance (q = 0.0263), as well as for higher macrophage abundance (q = 0.00162). Naive B cells were not significantly associated with survival (q = 1.00).

**Table 2.** Coefficients, statistics, p, and q-values for each immune cell term in a Cox proportional hazards model fit to predict overall patient survival, including diagnosis age, sex, cancer type, and number of years smoked as covariates.

|  | **Coefficient** | **exp(Coef.)** | **z-statistic** | **p-value** | **q-value** |
| --- | --- | --- | --- | --- | --- |
| **T cells** | -1.25217 | 0.285882 | -3.9169 | 8.97E-05 | 0.000538 |
| **Macrophages** | 0.694023 | 2.001752 | 3.460928 | 0.000538 | 0.001615 |
| **B cells** | -0.52134 | 0.593723 | -1.43318 | 0.151808 | 0.227711 |
| **NK cells** | -1.99242 | 0.136365 | -2.48022 | 0.01313 | 0.026261 |
| **Dendritic cells** | 0.18663 | 1.205181 | 0.368952 | 0.712164 | 0.712164 |
| **Misc. Myeloid** | 0.516746 | 1.676563 | 0.99876 | 0.317911 | 0.381493 |

We note that both the composition of the ITME and average patient age varies by cancer type. Therefore, to determine the variance in age-related effects that occur within different cancer types, we then evaluate the association between age and immune composition for each cancer type with at least 100 samples that could be successfully deconvolved by the MIXTURE algorithm (Supplemental Figure 3). We identify considerable heterogeneity in the effect of age on immune cell type abundance across cancer types, though macrophages generally increase in abundance, while T cells generally decrease (Figure 2A), consistent with the results of our pan-cancer analysis. Breast carcinomas have the most significant decrease in T cell abundance and the most significant increase in macrophage abundance with age overall.

**Figure 2–.**
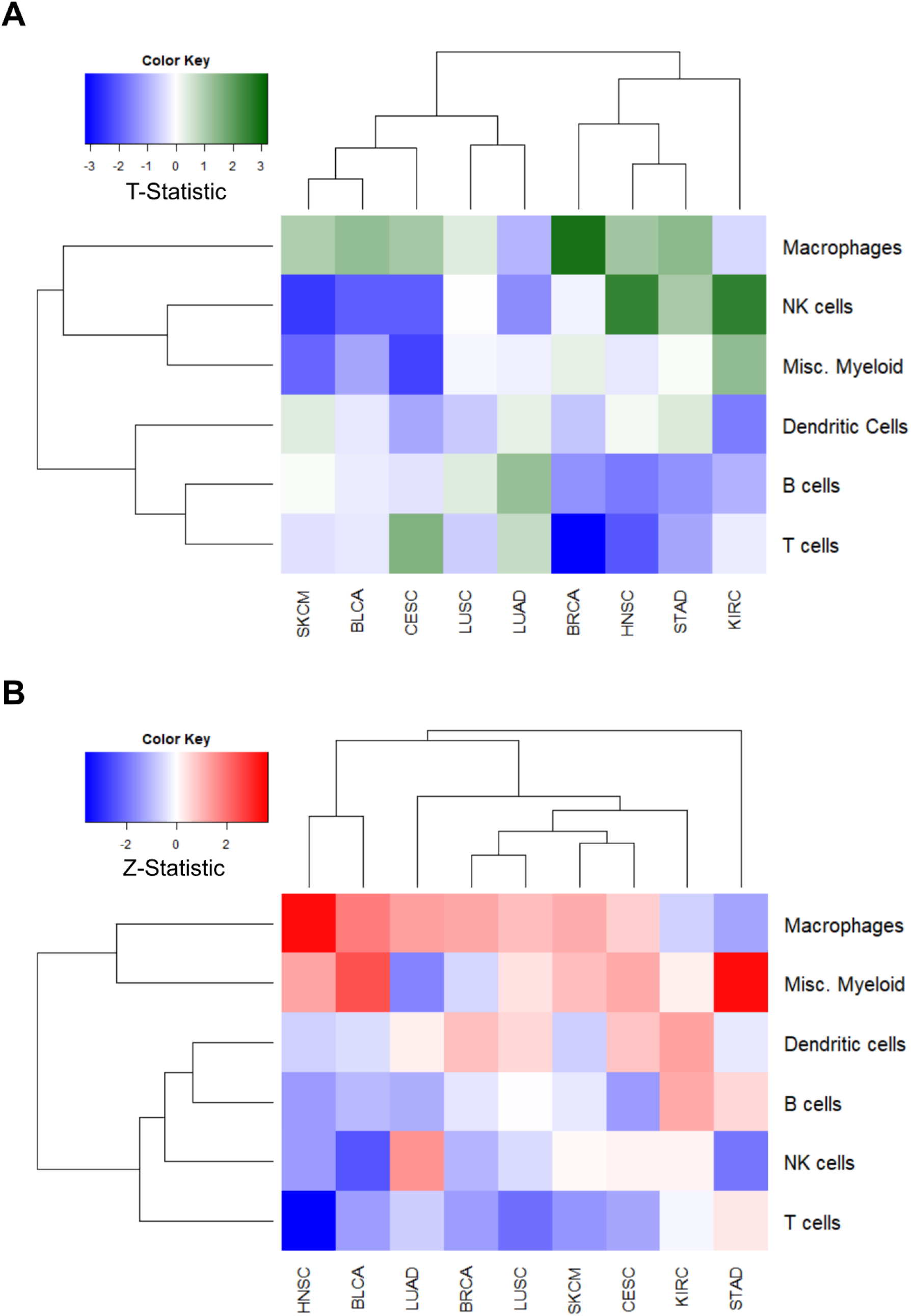
Age and survival association of immune cell types within cancer types. **A** Heatmap of the T-statistics from the covariate-adjusted linear model fit for the association of each immune cell type to age. Each square represents the significance of the diagnosis age term within a linear model for the labeled immune cell type, plotted for each cancer type in TCGA with at least 100 patients after filtering for significant immune deconvolution results. We observe that macrophages generally increase in abundance with age, while T cells and B cells generally decrease in abundance. **B** Heatmap of the covariate-adjusted Cox proportional hazards where each square represents the z-statistic for the survival prognostic of each immune cell type within each cancer type in TCGA with at least 100 patients after filtering. We observe that macrophages and other Myeloid cells are generally poor prognostics and that T cells are generally good prognostics.

To further distinguish the relative role of age and cellular composition of the ITME with patient outcomes, we then fit a Cox proportional hazards model for the effects of variation in each immune cell type across cancer types, including diagnosis age as a covariate (Figure 2B). Macrophages and other myeloid-derived cells are generally poor prognostics across cancer types, while T and NK cells are generally good prognostics. Head and neck squamous cell carcinomas (HNSC) have the most significant survival association with both macrophages and T cells. If HNSC cases are subdivided into HPV-negative and HPV-positive patients, this association only recapitulates among the HPV-negative cohort, emphasizing the importance of ITME composition for HPV-negative HNSC cases.

### Age-related immune changes in non-cancerous tissues differ from the observed shifts in the aging ITME among T cells, macrophages, B cells, and NK cells

To compare the effect of aging in the ITME to that on the immune cell compositions of normal tissues, we applied MIXTURE to GTEx consortium RNA-sequencing data of post-mortem samples from individuals without cancer (GTEx Consortium et al., 2017) to infer cell type abundance across tissues. These results provide a non-cancer baseline for immune changes that occur across many individuals of varying ages to compare with our observations from TCGA tumor data. Similar to the TCGA analysis, we fit a linear model to each cell type in order to determine associations between cell type abundance and age both across and within normal tissues.

In contrast to our findings in the pan-cancer ITME, in pan-tissue analyses we observe no significant change in overall T cell abundance with age (q = 0.565) (Figure 3A, Supplemental Table 2). We further fail to find significant changes in macrophage levels (q = 0.565) with age (Figure 3B). However, we do observe decreases in overall B cell (q = 3.51 x 10^-4^) (Figure 3C) proportion and increases in NK cell proportion (q = 1.07 x 10^-14^) (Figure 3D). This result demonstrates that there are considerable differences between the effects of aging on the abundance of immune cells in non-cancer tissues and the ITME.

**Figure 3–.**
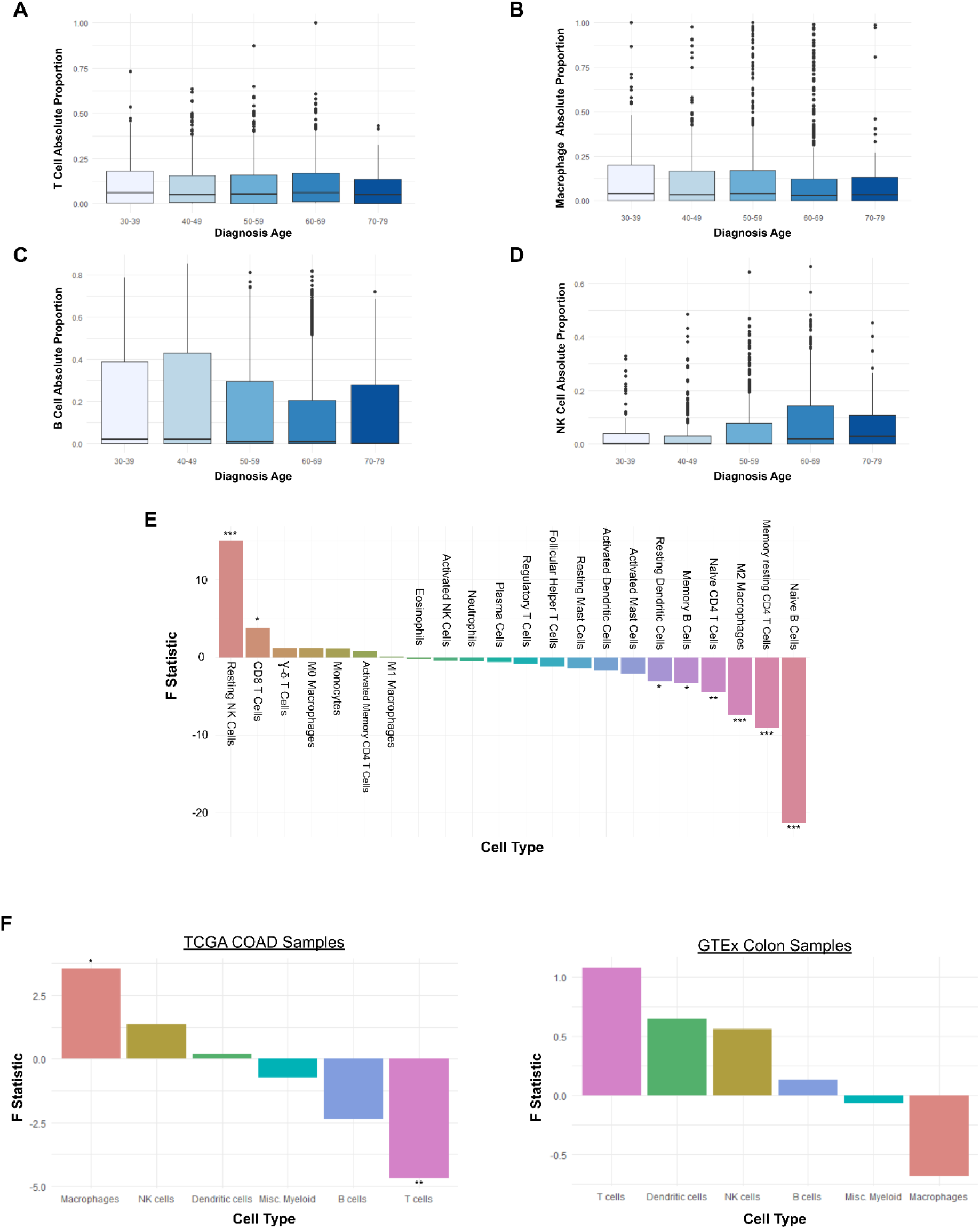
Immune changes with age in healthy tissues GTEx immune cell changes with age. **A-D** Immune abundance with age in GTEx tissues for T cells, Macrophages, B cells, and NK cells, respectively. **E** Age associations of 22 immune subtypes in GTEx tissues. Plotted by the F statistic for an ANOVA across age groups. **F** Comparison of F statistics for ANOVA across age groups for TCGA Colon Adenocarcinomas (COAD) against GTEx Colon samples. Opposite trends with age are observed for macrophages and T cells in this tissue. * indicates a p-value less than 0.05 and ** 0.01.

Across the 22 immune cell subtypes from MIXTURE, we find that naive B cell abundance is significantly decreased in GTEx samples with increasing age, consistent with our findings in the ITME (q-value of 9.07 x 10^-16^ and 0.0305, respectively). However, among GTEx data, we also observe significant decreases in memory CD4 T cells (q = 3.30 x 10^-7^), naive CD4 T cells (q = 0.00604), M2 macrophages (q = 3.59 x 10^-5^), memory B cells (q = 0.0315), and resting dendritic cells (q = 0.0427), as well as significant increases in abundance of resting NK cells (q = 5.33 x 10^-11^) and CD8 T cells (q = 0.0157) (Figure 3E). These findings represent clear departures from what we observe within the ITME, suggesting that tumors interact with the aging immune system to selectively prevent or increase infiltration of certain immune cell types that exist systemically, in a way that differs from the tumor immunosuppression that occurs in younger patients.

To directly compare normal age-related immune changes in a particular tissue to those that occur in tumors within the same tissue, we individually assay the immune associations with age for colon, lung, and breast tumors in TCGA with those found for all normal colon, lung, and breast samples from GTEx (Figure 3F, Supplemental Figure 4). Consistent with what we find pan-cancer and pan-tissues, we observe an increase in macrophage infiltration and a decrease in T cell infiltration among colorectal adenocarcinoma patients, while there is no significant change observed for colon samples from GTEx, and the non-significant trend that we do observe is reversed for both cell types. Similar immune cell type differences are found between cancerous and non-cancerous tissues in lung and breast samples. These results demonstrate that systemic age-related immune changes cannot be assumed to translate to the ITME. Particularly, they suggest that an interaction occurs between the phenotype of aging immune cells and the immunosuppressive signaling of cancers that generally increases the infiltration of macrophages and decreases the infiltration of T cells and NK cells in older patients.

### Systemic age-related increases in M1/M2 macrophage ratio and decreases in CD8/CD4 T cell ratio in non-cancerous tissues do not recapitulate in the ITME

We hypothesize that the tumor-specific changes to the ITME with age are associated with the modifications that cause tumors to evade immune attack. A higher ratio of M1/M2 macrophages has been previously found to be a positive survival prognostic (Chanmee et al., 2014), which we evaluate using a Cox proportional hazards model for overall survival in TCGA. We find that an increased M1/M2 ratio is a generally good prognostic pan-cancer, including age and cancer type as covariates (q = 0.0165). As the trends displayed in Figure 3E would suggest, the M1/M2 macrophage ratio significantly increases (q = 4.12 x 10^-4^) with age within normal tissues from GTEx data (Supplemental Figure 5A). By contrast, the M1/M2 tumor infiltration ratio does not change with age pan-cancer in tumor tissues in TCGA data (q = 0.314) (Supplemental Figure 5B), consistent with a signature of decreased immune activation relative to the rest of the body. Likewise, greater T cell killing would be expected to be associated with better prognosis. In TCGA, CD8/CD4 ratio has a minor favorable association with patient survival which falls short of 0.05 statistical significance (q = 0.0977). We observe an increase in the CD8/CD4 T cell ratio with age in GTEx samples (q = 2.87 x 10^-15^) (Supplemental Figure 5C) that does not recapitulate within the ITME (q = 0.314) (Supplemental Figure 5D).

### T cell receptor clonality decreases with age within the ITME while the number of tumor mutations increases

The overall decline in the total number of unique T cell receptor (TCR) clones with age (Yager et al., 2008), (Britanova et al., 2014), (Egorov et al., 2018) is well established in the literature. The process of thymic involution (the loss of thymus tissue with age) eventually ends the production of naive T cells and is the major driver of normal age-related decreases in T cell clonality (Aspinall and Andrew, 2000). To quantify aging-related changes in TCR clonality in the ITME, we leveraged results previously generated with the miTCR algorithm (Bolotin et al., 2013) by (Thorsson et al., 2018) to determine the association between TCR clonality and age. We define our metric of clonal diversity as the Shannon entropy multiplied by the number of unique clones divided by the total number of TCR sequencing reads (this correction is important because we have already established that T cell abundance decreases with age and we wish to correct this metric for the expected lower number of T cells to be sequenced among the samples from older patients). We determine that this TCR clonality measure significantly decreases with age for pan-cancer TCGA samples, including cancer type as a covariate (p = 1.48 x 10^-8^) (Table 3, Supplemental Figure 6A). We then fit a Cox proportional hazards model with age and cancer type as covariates and determine that decreased TCR clonality is a significant negative prognostic for overall survival (p = 3.34 x 10^-5^) (Table 3). This result suggests that the reduced ability to recognize antigens in older individuals leads to reduced T cell killing of tumor cells and hence worse outcomes.

**Table 3.** Coefficients, statistics, and p-values for the age term of the linear model for both TCR clonality and number of tumor mutations in TCGA data, with cancer type included as a covariate. These values are also listed for the TCR clonality term and the log number of tumor mutations term from a Cox proportional hazards model fit to overall survival with diagnosis age and cancer type included as covariates.

|  |  | **Coefficient** | **Statistic** | **p-value** |
| --- | --- | --- | --- | --- |
| **TCR Clonality** | **Diagnosis Age** | -0.00509 | -5.6694 | 1.48E-08 |
|  | **Cox Hazards** | -0.08812 | -4.14902 | 3.34E-05 |
| **Log Tumor Mutational Burden** | **Diagnosis Age** | 0.0102449 | 13.454 | 6.88E-41 |
|  | **Cox Hazards** | -0.053340 | -2.542 | 0.011 |

However, a related consideration is the increasing accumulation of mutations known to occur with age (Tomasetti et al., 2017) and the potential accompanying increase in immunogenic mutations. Higher tumor mutational burden has been shown to correlate with improved outcomes in ICB immunotherapy (Yarchoan et al., 2017), (Goodman et al., 2017), which suggests the possibility that the increasing number of mutations accumulated with age may in some cancers offset the loss of TCR diversity. As has been previously reported (Chalmers et al., 2017), (Qing et al., 2020), we find that tumor mutational burden significantly increases with age (p < 1×10^-16^) (Supplemental Figure 6B) pan-cancer in TCGA, including cancer type as a covariate (Table 3). This trend recapitulates within most cancer types, though lung adenocarcinomas and uterine carcinomas are notable exceptions (Supplemental Figure 6C). High mutational burden among younger patients with lung cancer is likely due to the highly mutagenic effects of cigarette smoking, while the highly mutated uterine tumors are likely the result of a hypermutated subset previously discovered among the TCGA-UCEC cohort (Cancer Genome Atlas Research Network et al., 2013). A higher tumor mutational burden is a positive survival prognostic pan-TCGA, as determined by a Cox proportional hazards model, including age and cancer type as covariates (p = 0.011) (Table 3). Thus, while the ability to recognize antigens may decrease with age, the space of tumor antigens for a given TCR to match with will likely increase, presumably at least partially offsetting the decreased T cell capacity to recognize tumors caused by loss of TCR diversity.

### Age-related increases in tumor mutational burden associates with promoter demethylation and increased expression of immune checkpoint genes in tumor samples

The ability of TCR sequences to recognize tumor antigens is of particular clinical relevance in the context of ICB immunotherapy. Another important factor for the efficacy of ICB immunotherapy is the expression of target genes and their complementary receptors such as *PD1*, *PDL1, CTLA4, CD80,* and *CD86* (Taube, 2014). We therefore performed differential expression analysis for these genes with age in both TCGA tumor samples and normal GTEx tissue samples, including cancer type and tissue type as respective covariates. Among TCGA samples, *CD80* and *PDL1* expression increases with age (q-values 0.0116 and 0.0299), while no detectable expression changes occur with age for *CTLA4, PD1,* and *CD86* (q-values of 0.779, 0.693, and 0.0834) (Table 4), suggesting that older patients display increased tumor cell expression of these immune checkpoint genes. Another possibility is that this change occurs due to the increased number of infiltrating macrophages we have shown accumulate with age, as PDL1 and CD80 can be expressed on macrophages as well (Hartley et al., 2018). We further investigate age-related changes in expression of these genes within each cancer type (Supplemental Figure 7). There is considerable heterogeneity in the effect of age on the expression of these genes, though *PDL1, CD80,* and *CD86* are more likely to increase in expression with age, while *PD1* and *CTLA4* expression is more variable, suggesting that tumor expression of immune checkpoint genes is more affected by age than tumor infiltrating T cell expression of immune checkpoint genes.

**Table 4.** Differential expression results for immune checkpoint genes involved in currently available ICB immunotherapies. The results are shown for the association with age, both including tumor mutational burden as a covariate and without.

|  | Gene | LogFC | t-statistic | p-value | q-value |
| --- | --- | --- | --- | --- | --- |
| **Age +**<br>**Cancer Type**<br>**Adjusted** | **CD80** | 0.00361 | 3.002352 | 0.002686 | 0.011553 |
|  | **CD274** | 0.003053 | 2.612337 | 0.009007 | 0.029899 |
|  | **CD86** | 0.002146 | 2.130932 | 0.03312 | 0.083376 |
|  | **CTLA4** | -0.00084 | -0.5747 | 0.56551 | 0.69286 |
|  | **PDCD1** | -0.00062 | -0.42127 | 0.67357 | 0.778683 |
| **+**<br>**Mutational**<br>**Burden**<br>**Adjusted** | **CD80** | 0.002983 | -0.90527 | 2.38488 | 0.017105 |
|  | **CD86** | 0.002127 | 2.9119 | 2.035515 | 0.041828 |
|  | **CD274** | 0.00198 | 2.124811 | 1.618133 | 0.105669 |
|  | **CTLA4** | -0.00204 | 0.12074 | -1.34813 | 0.177652 |
|  | **PDCD1** | -0.002 | 0.334003 | -1.30557 | 0.191731 |

Among the non-cancerous samples from GTEx, *PD1* and *PDL1* expression significantly increases with age (q-values of 2.93 x 10^-7^ and 0.00166), *CD86* expression does not significantly change (q = 0.891), and there was no data available for *CTLA4* or *CD80* (Supplemental Data). The increase in *PD1* expression is possibly due to the increased presence of exhausted CD8 T cells, which have been previously reported to accumulate with age and express both *PD1* and *TIM3* (Lee et al., 2016), the latter of which is also expressed at increased levels with advanced age in GTEx data (q = 2.157 x 10^-5^). The increased inflammation observed in older individuals (Fulop et al., 2017), (Kovtonyuk et al., 2016) may explain the increased expression of *PDL1* in older individuals from GTEx. Among the inflammatory pathways up with age, we particularly note that the GO term for response to interferon gamma significantly increases in GTEx (q = 0.00297), which has been reported to stimulate *PDL1* expression (Flies and Chen, 2007).

Tumor mutational burden is an important clinical predictor of response to immunotherapy that increases with patient age. To determine if these age-related changes in immune checkpoint gene expression are associated with the increase in tumor mutational burden that we described previously (Supplementary Figure 6B), we included it as a covariate and repeated the differential expression analysis. *PDL1* expression no longer appeared significantly associated with age (q = 0.207), while *CD80* expression became borderline (q = 0.0502) (Supplemental Data). These data indicate that increased expression of *PDL1* with age most directly associates with age-related increases in tumor mutational burden.

Due to previous work suggesting that DNA methylation regulates tumor expression of *PDL1* (Asgarova et al., 2018), (Micevic et al., 2019), we hypothesize that the observed expression increases of immune checkpoint genes are largely driven by epigenetic changes. We leverage merged 450k and 27k methylation array data from TCGA (Thorsson et al., 2018) and find that methylation of CpGs annotated to the *PDL1* promoter region significantly decreases with age pan-cancer (q = 3.27 x 10^-10^), including cancer type as a covariate. Methylation of *CTLA4* and *CD80* annotated CpGs also decrease with age (q values of 4.33 x 10^-4^ and 6.01 x 10^-5^) (Table 4). These results suggest that DNA methylation changes lead to the observed expression increases in *PDL1* and *CD80* with increasing age and that age-related increases in tumor mutational burden promote selective pressure for epigenetically mediated up-regulation of *PDL1*.

### Corresponding shifts in DNA methylation and expression in TCGA samples suggest profound changes in the aging tumor microenvironment

In order to further investigate the role of age-related methylation changes on the ITME, we identified all genes annotated to have some immune role by InnateDB (Breuer et al., 2013) within TCGA expression and methylation data. In order to find immune genes that are regulated by age-related methylation changes at promoters of these genes, we then selected the following two sets of differentially expressed genes: those that had significant increases in expression and significant decreases in methylation with age, and those with significant decreases in expression and significant increases in methylation with age. We note that increased methylation of annotated promoter CpGs was much more likely to indicate that the corresponding gene would decrease in expression (218 anti-correlated vs 66 correlated with age) than decreased methylation was to indicate that a gene’s expression would increase (113 correlated vs 94 anti-correlated with age) (Supplemental Table 3).

Among the genes with increased expression and decreased methylation in TCGA, we find the GO immune regulation term (q = 5.90 x 10^-25^), the regulation of T cell activation term (q = 2.93 x 10^-5^), and innate immune response term (q = 1.74 x 10^-11^) significantly enriched (Figure 4A) (the full list of GO enrichments is available in Supplemental Data). Of particular note is the connected group of T cell regulatory genes *PDL1, CD80, LAG3, HAVRC2,* and *IL10. LAG3* has been shown to be of importance in immune infiltration and overall survival in renal cell carcinoma (Klümper et al., 2020), acts as an important player in intratumoral T cell exhaustion in lymphoma (Yang et al., 2017), and has been suggested as a potential therapeutic target for new ICB strategies (Long et al., 2018). *HAVRC2* (also known as *TIM3)* has been shown to be an important inhibitory T cell receptor, as well as a defining characteristic of exhausted T cells in concert with *PD1* (Lee et al., 2016), (Wolf et al., 2020). *IL10* has been shown to be a direct inhibitor of CD8 T cell function (Smith et al., 2018). We thus observe a set of methylation-associated expression changes with increasing age that are expected to have a considerable detrimental effect on T cell tumor killing.

**Figure 4–.**
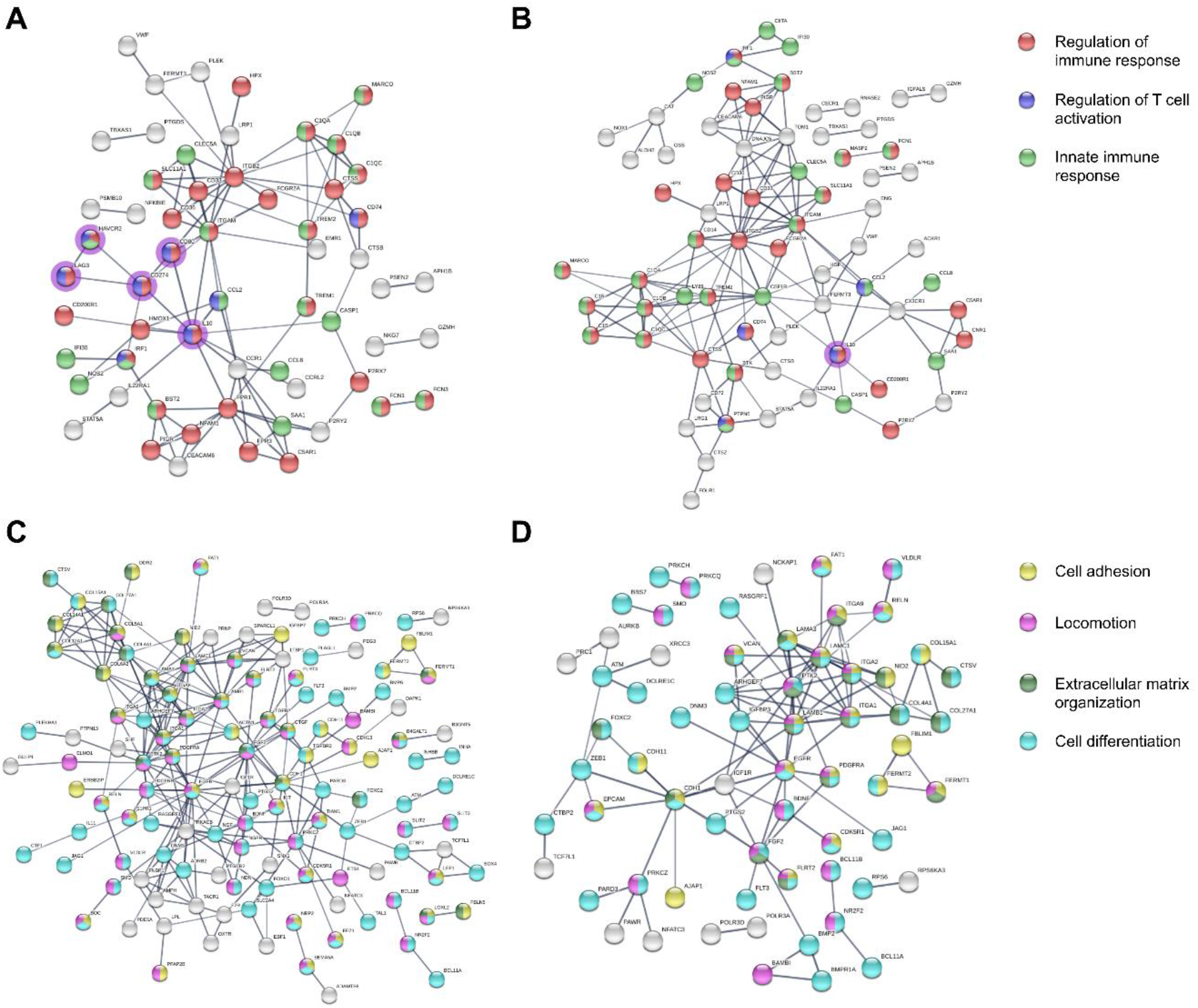
Corresponding methylation and expression changes with age. STRING Database queries for all genes with significantly **A** increased expression and decreased methylation with age, where genes that have previously been strongly associated with decreased T cell cytotoxicity have a purple halo **B** increased expression and decreased methylation with age, adjusting for tumor mutation count, where genes that have previously been strongly associated with decreased T cell cytotoxicity have a purple halo **C** decreased expression and increased methylation with age, **D** decreased expression and increased methylation with age, adjusting for tumor mutation count.

Next, we investigated if these methylation and expression changes are related to the age-associated TMB changes, like we observed for *PDL1* and *CD80.* We performed the same analysis of immune-related genes in InnateDB, adjusting for tumor mutational burden for both differential expression and methylation, and only *IL10* retains significantly increased expression and decreased methylation of the aforementioned five genes (Figure 4B). Therefore, the effect of age on T cell activation in our model seems to be largely explained by age-related changes in tumor mutational burden, while other age-related factors have a larger impact on the increased expression of a number of genes related predominantly to innate immunity. For example, another 9 genes annotated to the innate immune response GO term are significant for the TMB corrected analysis (25 vs 34 total).

The genes with increased CpG methylation and decreased expression with age in TCGA samples are significantly enriched for a number of GO terms, notably including cell adhesion (q = 9.38 x 10^-44^), cell differentiation (q = 5.83 x 10^-31^), locomotion (q = 1.84 x 10^-21^), and extracellular matrix organization (q = 6.18 x 10^-21^) (Figure 4C) (full list provided in Supplemental Data). This gene set also includes several growth factors and receptors with major known roles in tumor biology, such as *EGFR* (Normanno et al., 2006), *FGF2* (Soufla et al., 2005), *PDGFRA* (Velghe et al., 2014), and *IGF1R* (Pandini et al., 1999). All of these terms and genes remain significant after correcting for tumor mutational burden (Figure 4D), though major transcription factors such as *SOX4* and *FOXO1* drop out, as do the growth factors *TGFB2* and *NGF*, indicating that high mutational burden may play a role in age-related gene methylation for a subset of these genes. However, most of these gene expression and methylation changes appear to be mediated through other age-related effects. Epigenetic silencing of these pathways and genes would be expected to have a considerable effect on the tumor microenvironment, as they play major roles in critical tumor processes such as growth, metastasis, dedifferentiation, and angiogenesis.

### GO gene set enrichment differences between TCGA and GTEx data with age suggest interactions of tumor signaling and aging biology produce substantial changes in tumor cellular processes and microenvironment

To further investigate possible causes for the observed differences in immune infiltration between aged tumors and healthy aged tissues, we compare GO term enrichment based on gene expression changes with age from TCGA and GTEx samples. We primarily investigate four sets of GO terms: those up in both TCGA and GTEx with age, those up in TCGA and down in GTEx, those up in GTEx and down in TCGA, and those down in both TCGA and GTEx (Supplemental Figure 8) (full list of GO terms provided in Supplemental Data). The pathways up in both are highly immune related, indicating an increase in inflammation with age that has previously been termed inflammaging (Fulop et al., 2017), (Franceschi et al., 2000). The increases in antigen binding, MHC complex, and Interferon gamma signaling terms are of relevance to T cell recognition and activity. Of particular note may be the increase in Interferon gamma signaling, which is known have a role upregulating *PDL1* expression (Flies and Chen, 2007), and thus may be involved in the expression increase of *PDL1* with age, along with the previously noted relation to tumor mutational burden. Among those pathways up in TCGA and down in GTEx, the Mitochondrion term seems the most potentially relevant, as energetics is a major factor in tumor biology, and mitochondria additionally play a considerable role in regulating cell death pathways as well as immune regulation (Weinberg et al., 2015), (Breda et al., 2019). Regulation of proliferation, positive regulation of development, locomotion, and biological adhesion are up in GTEx and down in TCGA, with potential relevance for tumor growth and metastasis. These terms match closely with the terms found to be silenced by DNA methylation with age in the previous section, indicating that these are cancer specific age-related effects as well. There thus appears to be another interaction of aging biology and tumor signaling that may alter the development of several major hallmarks of cancer in older patients. Finally, cell cycle terms are down in both cohorts with increasing age. In the context of the altered age-related immune infiltration in tumors compared to normals, these data demonstrate that there are inflammation increases with age in both tumor and healthy aged tissues. Thus, the observed changes in immune cell abundance between the tumors of old patients and the tissues of old individuals are likely the result of an interaction between tumor immunosuppressive signaling and the altered phenotype of aged immune cells.

## Discussion

We demonstrate that patient age associates with several changes in ITME composition that do not occur in non-cancerous tissues. This result indicates that tumor signaling interacts with the aging immune system and/or environment to modulate immune infiltration and produce a more immunosuppressive microenvironment in older patients than would be predicted by normal immune aging in isolation. We further investigate the complex interplay of age-related shifts in TCR clonality, tumor mutational burden, and T cell exhaustion that appear to modulate the demethylation and increased expression of immune checkpoint genes in the tumors of older patients.

In contrast to our analysis of ITME cell type composition shifts with age, there is an existing literature on normal age-related immune changes, which allows for validation of the results of the MIXTURE method in this work with an established base of knowledge. The observed decreases in overall T cell abundance within tumors did not recapitulate in normals from GTEx and have not been reported to occur systemically. While this may seem a strange phenomenon given the process of thymic involution, it has been previously found that it is mostly naive T cells that decrease in abundance with age, while the population of memory T cells proportionally increases (with accumulation of exhausted, less functional effector T cells as individuals reach very advanced age) (Hulstaert et al., 1994), (Alpert et al., 2019). Thus, overall T cell abundance remains relatively constant, as we observe in our analysis of GTEx data. This ITME specific departure may have implications for T cell based immunotherapies in elderly patients, suggesting the possibility of targeting the as of yet unknown mechanism repressing infiltration to potentially improve therapeutic efficacy in older patients.

We observe increased infiltration of macrophages in tumor samples, without a particular bias in M1/M2 polarization, while in normal GTEx data we identify no change in macrophage abundance overall, but find that the M1/M2 ratio increases. Previous studies have suggested that old age alters macrophage polarization such that the same signals that would produce polarization in younger individuals do not produce differentiated macrophages with all the characteristics generally ascribed to M1 and M2 subtypes (Mahbub et al., 2012). Thus, these results must be interpreted carefully, but an increase in tumor infiltrating macrophages still appears to be a poor survival prognostic in older individuals, which suggests that therapeutics being developed to target the generally negative effects of tumor-associated macrophages (Chanmee et al., 2014), (Poh and Ernst, 2018), (Lee et al., 2019) may be of particular impact in patients of advanced age.

A further observation of note is that we find a significant increase in NK cell abundance in GTEx, which we do not identify in TCGA tumor data. It has been previously shown that overall NK cell abundance increases with age, though their cytotoxicity was diminished relative to younger controls (Gounder et al., 2018). It therefore seems that NK cell tumor infiltration is inhibited with increasing patient age because this general increase in abundance is not reflected in the ITME. The role of NK cells in cancer immunity is gaining increasing appreciation (Freeman et al., 2019), (Chiossone et al., 2018), with some evidence to suggest an important role in emerging immunotherapies as well (Shimasaki et al., 2020).

Systemic loss of TCR clonality with age (Britanova et al., 2014) is reflected in the ITME, which, along with the decreased T cell infiltration we identify, raises questions about the efficacy of immunotherapies in patients of advanced age due to the value of TCR diversity as a biomarker to identify patients who will benefit from ICB immunotherapy (McNeel, 2016). The possibility of reduced efficacy has been partially addressed by previous studies. There are somewhat mixed results as to the benefit of ICB for patients of advanced age, though most studies and metaanalyses of available clinical trial data suggest patients experience no reduced benefit (Kugel et al., 2018), (Elias et al., 2018), (Daste et al., 2017), (Jain et al., 2019). The observed increases in tumor mutational burden with age may explain these results at least partially, as these additional mutations provide a larger space of antigens for the reduced number of unique TCRs to recognize. We further find that the tumor mutational burden increases with age appear to mediate a decrease in methylation and increase in expression of PDL1, though we cannot rule out that age-related increases in Interferon gamma signaling play a role as well. From these results, we hypothesize the causal model outlined in Supplemental Figure 10, whereby TCR Diversity, Tumor mutational burden, and T cell infiltration changes with age together mediate the cytotoxic capacity of T cells on tumors and thus the selective pressure for the expression of immune checkpoint genes. Given the results of the aforementioned meta-analyses, this model suggests that increased tumor mutational burden (and the corresponding increase in PDL1 expression) is largely able to compensate for the loss of T cell infiltration and TCR diversity such that older patients end up doing roughly as well on ICB immunotherapy as their younger counterparts despite several characteristics that are generally disadvantageous, though unaccounted factors may be at play as well. Currently, ICB is still used less often among elderly patients (Jain et al., 2019), (Hurez et al., 2018) due to concerns about efficacy and safety. Our results indicate that older individuals express increased levels of PDL1, which likely allows for the equal level of response to that of younger patients. This suggests that ICB use for the elderly should be further investigated and likely expanded. Additionally, our model suggests that if T cell infiltration into the tumors of older patients can be increased, they might respond even better to ICB immunotherapy than young patients do.

We identify a set of genes that are hypermethylated and lower-expressed with increasing age. These genes are significantly associated with growth, metastasis, and angiogenesis, which are considered to be some of the hallmarks of cancer (Hanahan and Weinberg, 2011). We further find that these terms are not enriched among GTEx samples with age, demonstrating another interaction between aging biology and tumor biology that produces an age-related tumor phenotype that is distinct from the age-related phenotype in normal tissues. Within this set of genes we observe what appears to be DNA methylation-mediated silencing of genes associated with cell differentiation, which previous studies (Widschwendter et al., 2007), (Easwaran et al., 2012), (Ohm and Baylin, 2007), (Schlesinger et al., 2007) indicate represents a tumor-specific hypermethylation of genes that locks the constituent cells into a malignant stem-like state. These changes appear to be more common in the tumors of elderly patients, suggesting epigenetic therapies may be of particular value in this population.

Given the age-related differences in immune infiltration observed, we identified both the differences and similarities in GO term enrichment with age for TCGA and GTEx samples. Inflammation and immune terms are up in both, including interferon gamma signaling, which likely has some impact on the observed increase in PDL1 expression (Flies and Chen, 2007). The differences in enrichment, however, are largely not directly immune related and do not ostensibly explain the observed differences in immune infiltration. However, some of these pathway differences, such as increased mitochondrial activity in tumors with age and decreased activity in normal aged tissues, may merit future investigation due to the potential role these changes may play on cellular energetics and survival.

To determine if age-related tumor mutational burden increases mediate the immune infiltration changes we observe with age, we included it as a covariate in analyses of immune infiltration derived from MIXTURE. We determined none of our observations changed substantially (Supplemental Data), demonstrating that the observed differences are mediated through other age-related factors. Thus, we conclude the T cell infiltration decreases, macrophage infiltration increases, and NK infiltration stability despite an increasing abundance in the body overall is most likely attributable to the hypothesis that exhausted or senescent phenotypes develop in immune cells as individuals age and make the immune cells of older patients more sensitive to the immunosuppressive signaling that is produced by most tumors (Lu et al., 2019). Another possibility is that some other aspect of aging biology extrinsic to immune cells interacts with the dysregulated signaling from tumors to prevent immune infiltration of certain cell types. However, these investigations do not provide any particular evidence of what that actor might be and therefore the former hypothesis should be favored by virtue of parsimony. Additional investigation is needed to determine whether either of these proposed mechanisms drives our observations, but they suggest the possibility of therapeutics targeting the immune senescence phenotype or the interaction of said phenotype and tumor immunosuppressive signaling to improve outcomes for older cancer patients.

It is important to note the limitations of bulk expression data for the analysis of tumor-infiltrating immune cells. Possibly most importantly is the consideration of immune cell function and quality (e.g. the question of whether T cells are exhausted or non-functioning is difficult to address from bulk data). Further, immune cell type deconvolution of bulk data does not lend itself to as thorough an exploration of immune subtypes as at single-cell resolution. Therefore, future single-cell pan-cancer characterization from projects such as the Human Tumor Atlas Network (Rozenblatt-Rosen et al., 2020) and normal tissue through the Human Cell Atlas are critical to further delineate the role that aging-related changes to immune cell function play in cancer. Nonetheless, characterization of the impact aging has on the ITME from bulk data can be a significant aid in the informed treatment of elderly patients.

Overall, these results suggest that patients of advanced age may benefit from immune modulation that promotes infiltration of immune cells that are already present in their body to produce a more favorable environment for therapeutic response and survival. Additionally, our results, along with previous meta-analyses of clinical trials, suggest that ICB use in elderly patients merits further study and expansion. Finally, these findings indicate that to fully appreciate the tumor biology and treatment needs of older patients, specific study into the effects of age in cancer is necessary, as we cannot necessarily rely on studies of aging in general to accurately describe the effects of age in tumors.

## Methods

### RNA-Sequencing Data

TCGA RNA-sequencing data processed and normalized according to https://docs.gdc.cancer.gov/Data/Bioinformatics_Pipelines/Expression_mRNA_Pipeline/ was downloaded from the GDC Data Portal on August 8th, 2019, filtering for all TCGA samples with patients above 30 years of age. Patients under 30 were excluded to focus on ITME changes in adult populations, which are more likely to generalize to the majority of cancer patients.

GTEx RNA-sequencing counts version 8 were downloaded from the GTEx Portal on November 12th, 2019. Only individuals over 30 were included in the final analysis, to be comparable with filtering of TCGA. Characteristics of these cohorts are listed in Supplemental Table 4.

### Immune Cell Type Deconvolution from Bulk RNA-Sequencing Data

The MIXTURE algorithm (Fernandez et al., 2019) builds on the nu-Support Vector Regression framework used by CIBERSORT (Newman et al., 2015) for particular use with noisy tumor samples. MIXTURE applies Recursive Feature Selection to make the cell type deconvolution more robust to noise and collinearity, and was thus designed to improve performance on tumor data.

We run MIXTURE using a population-based null distribution and the nu-SVM Robust RFE method on the preprocessed RNA-sequencing data from both TCGA and GTEx. A signature expression matrix (LM22 from Newman et al) (Newman et al., 2015) is used to determine the proportion of 22 immune cell types in each sample. MIXTURE returns both relative and absolute proportions of immune cells. Absolute proportions were used for all analyses of TCGA and GTEx datasets. MIXTURE provides a p-value for the cell type deconvolution performed. Only samples with a deconvolution p-value less than 0.05 were used in the final analyses, leaving 3576 patient samples remaining in TCGA and 1689 in GTEx. A further 29 TCGA patients had received treatment prior to sample collection, and were removed to avoid biasing of results.

### Modeling the Association of Immune Cell Type with Age and Survival

Linear models are fit to investigate the association between the absolute proportion of each immune cell type and the initial diagnosis age in TCGA. The models are fit separately for each cancer type as well as jointly with cancer type and patient sex as covariates. Significance is assessed using Benjamini-Hochberg FDR correction for multiple testing across all cell types tested.

Higher order cell types are defined by adding together individual substituent cell type values and dividing by the sum of all cell types, the result of which is used as the predictor variable in the linear model (which immune subtypes correspond to which higher order cell types is shown in Supplemental Table 1).

In addition to the linear models, we made box and violin plots of the immune cell type absolute proportion by age group without additional covariate adjustment to visualize immune changes with age.

The relationships between overall survival and immune cell types are assessed using a Cox proportional hazards model fit with the survival R package Version 3.1-8. The model uses months of overall survival as the outcome and includes diagnosis age and sex as covariates. It is further stratified by cancer type.

GTEx data only provides the age group of each individual rather than the particular year of age at the time of sample collection, so an ANOVA is performed between absolute immune cell proportion and age group, including age and tissue type as covariates.

### Modeling the Age Associations of Normalized TCR Clonality and Number of Tumor Mutations

TCR clonality is assessed using miTCR (Bolotin et al., 2013) results previously published by Thorsson et al., 2018 (Thorsson et al., 2018). Our previous results demonstrated decreased infiltration of T cells with increasing age, so to avoid biasing our results, the Shannon Entropy is multiplied by the number of unique TCR clones divided by the total number of TCR reads. We then fit a linear model for the association of age with this TCR clonality measure, including patient sex and cancer type as covariates. We again use a Cox Proportional hazards model to assess if normalized TCR clonality is a relevant survival prognostic, using the same survival function and covariates as described above.

To find the association of age and number of tumor mutations we downloaded the mutation count provided for each sample from the GDC data portal. We log transformed the data due to the skewed distribution of tumor mutation counts and fit the values to a linear model and Cox proportional hazards model, using the same covariates as above.

### Differential Expression Analysis with Age

Differential expression analyses from both TCGA and GTEx data were performed on all samples from individuals of at least 30 years of age. The edgeR package Version 3.26.8 was used for normalization and identifying differentially expressed genes with age. Diagnosis age was modeled as a continuous variable, including cancer type as a covariate for the TCGA analysis and tissue type as a covariate for the GTEx analysis. Genes were considered differentially expressed below an FDR adjusted p-value of 0.05. A further differential expression analysis was performed on TCGA data, including mutation count as an additional covariate. Finally, a differential expression analysis for diagnosis age was performed on each cancer type separately that had at least 100 samples.

### Differential Methylation Analysis with Age

Merged 450k and 27k DNA methylation array data preprocessed by (Thorsson et al., 2018) was downloaded from GDC at https://gdc.cancer.gov/about-data/publications/panimmune. A linear model for diagnosis age was fit to data from each CpG, including cancer type as a covariate. CpG methylation was considered significantly different with age if the FDR adjusted p-value for the diagnosis age term was less than 0.05. Annotations of CpG sites to gene promoters were retrieved from the IlluminaHumanMethylation27k.db R package Version 1.4.8. The same analysis was repeated, additionally including mutation count as a covariate in the model.

### Finding Epigenetically Regulated Immune Genes

Genes annotated to have an association with immune function were downloaded from InnateDB (Breuer et al., 2013), (Lynn et al., 2008). Differentially expressed genes were subset to those overlapping with the set annotated as immune-related. These sets were then further subset to those that were up-expressed with age and down-methylated, as well as those down-expressed with age and up-methylated, in accordance with the canonical understanding of the effects of DNA methylation on gene expression.

These sets were produced for both the analysis including and not including mutation count as a covariate. The resulting gene sets were visualized using the STRING database (Szklarczyk et al., 2019), which shows the known or predicted interactions that the corresponding proteins would be expected to have.

### Gene Set Enrichment Analysis

The fgsea R package version 1.10.1 (Sergushichev, 2016) was used to perform gene set enrichment analysis from differential expression results with age from TCGA and GTEx, produced as described above. GO terms were downloaded from MsigDB (Liberzon et al., 2011) using the msigdbr R package Version 7.0.1. GO enrichment was compared between the data sets, identifying those terms significantly up in both, up in one and down in the other, and down in both with age.

## Code Availability

All analysis code is available on GitHub at: https://github.com/rossinerbe/ImmuneAgingAnalysis

## Acknowledgements

The authors would like to thank Ashani T. Weeraratna for her input on aging immunity in cancer. The authors would also like to thank Mara R. Lanis for her helpful discussions on TCR diversity and immune checkpoint genes.

The results shown here are based upon data generated by the TCGA Research Network: https://www.cancer.gov/tcga and the The Genotype-Tissue Expression (GTEx) Project https://www.gtexportal.org/.

## Supplemental Figures

**Supplemental Figure 1.**
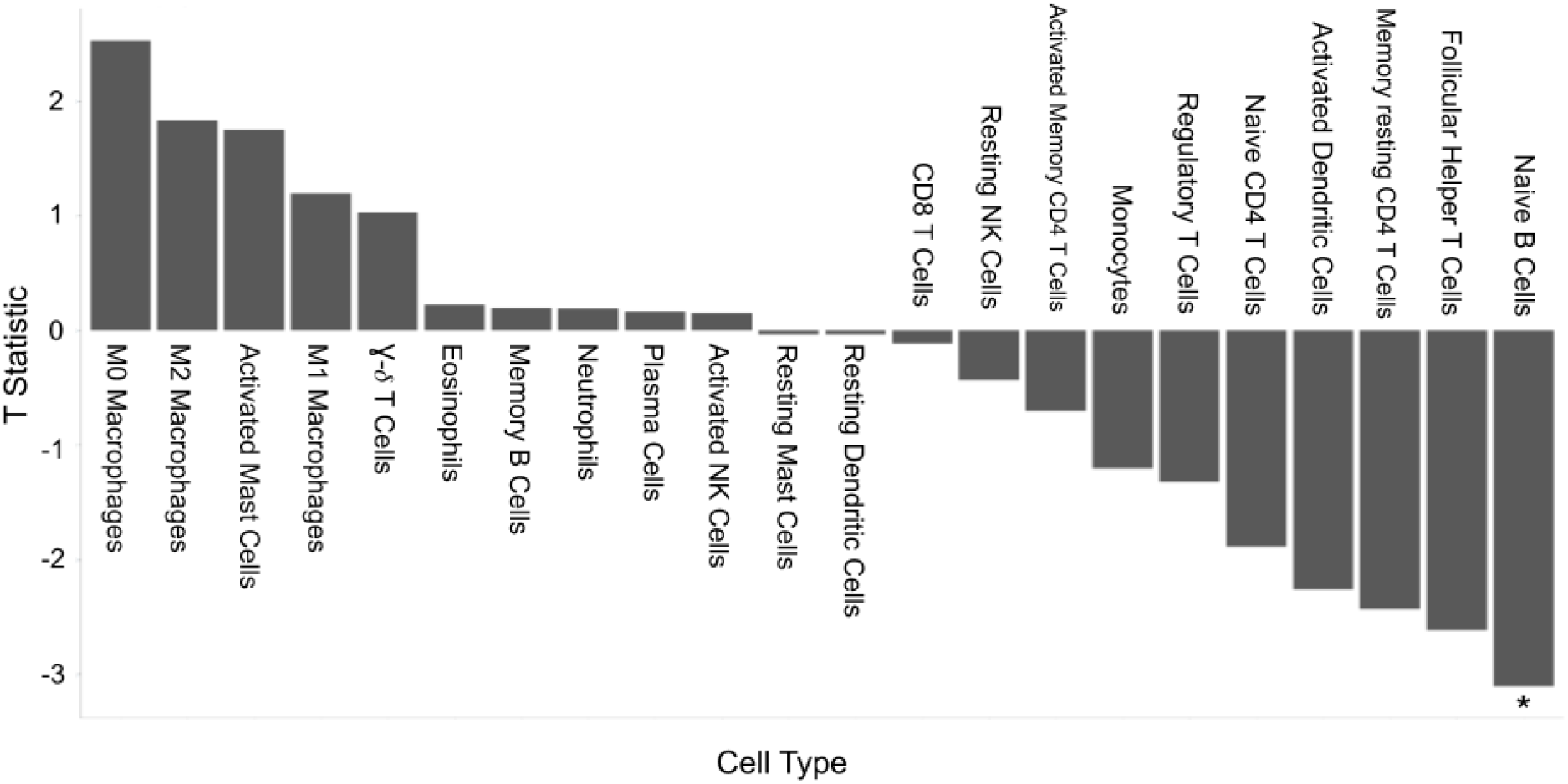
T statistics for the linear association of 22 immune cell subtypes with age pan-cancer within TCGA data. Naive B cells statistically significantly decrease with age. * indicates q-value less than 0.05.

**Supplemental Figure 2.**
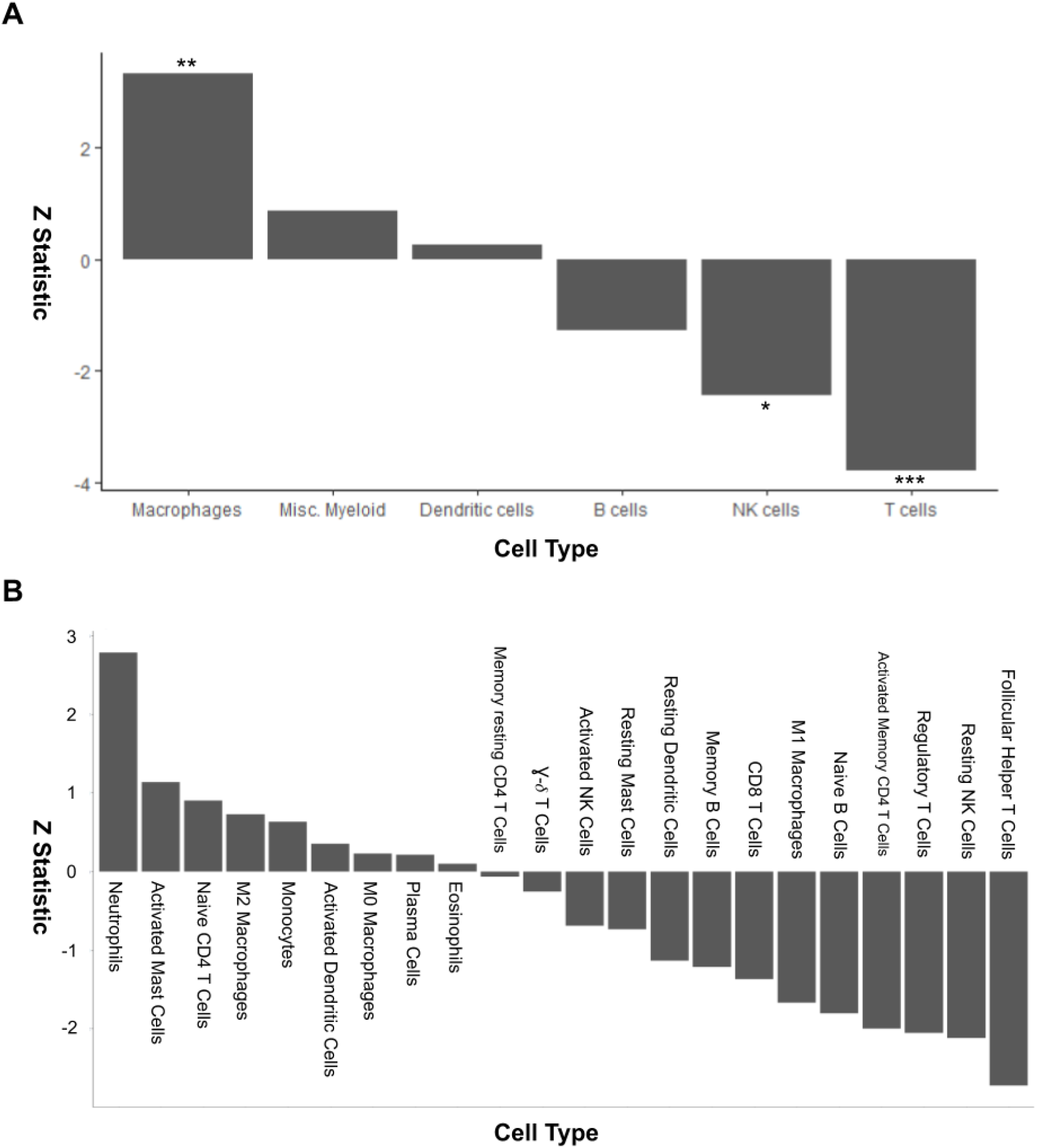
**A** Z-statistics for the survival associations of immune cell types pan-cancer as evaluated by a Cox proportional hazards model, including age, sex, cancer type, and smoking years as covariates. **B** Z-statistics for the survival associations of 22 immune cell subtypes as evaluated by a Cox proportional hazards model for overall patient survival pan-cancer including age, sex, cancer type, and smoking years as covariates. * indicates a p-value less than 0.05, ** 0.01, and *** 0.001.

**Supplemental Figure 3.**
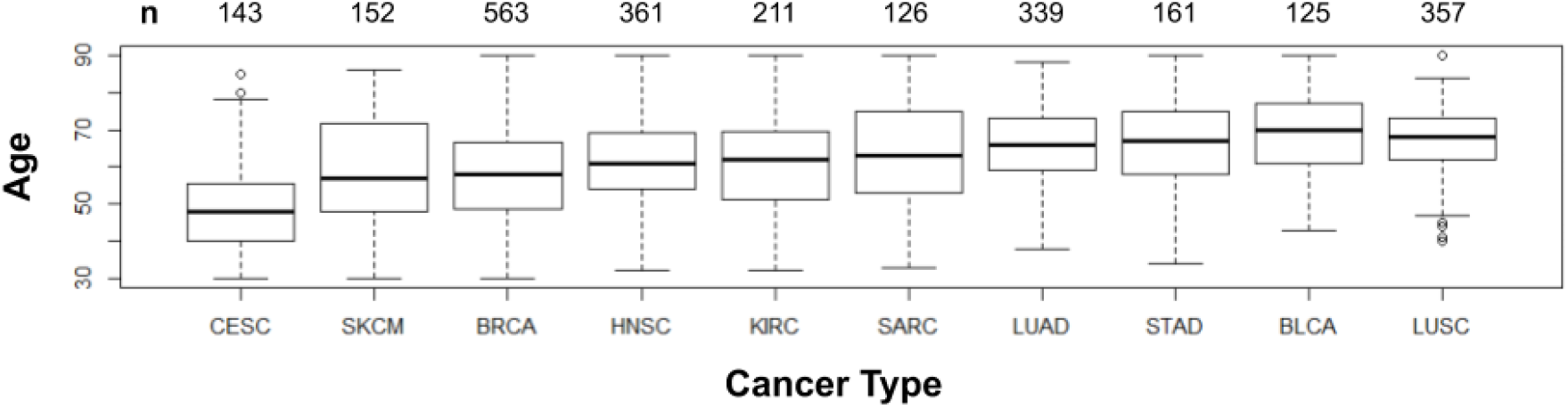
Boxplot of the age demographic for each of the cancer types with at least 100 observations that were successfully deconvolved by immune cell type. The number of patients with each cancer is plotted above.

**Supplemental Figure 4.**
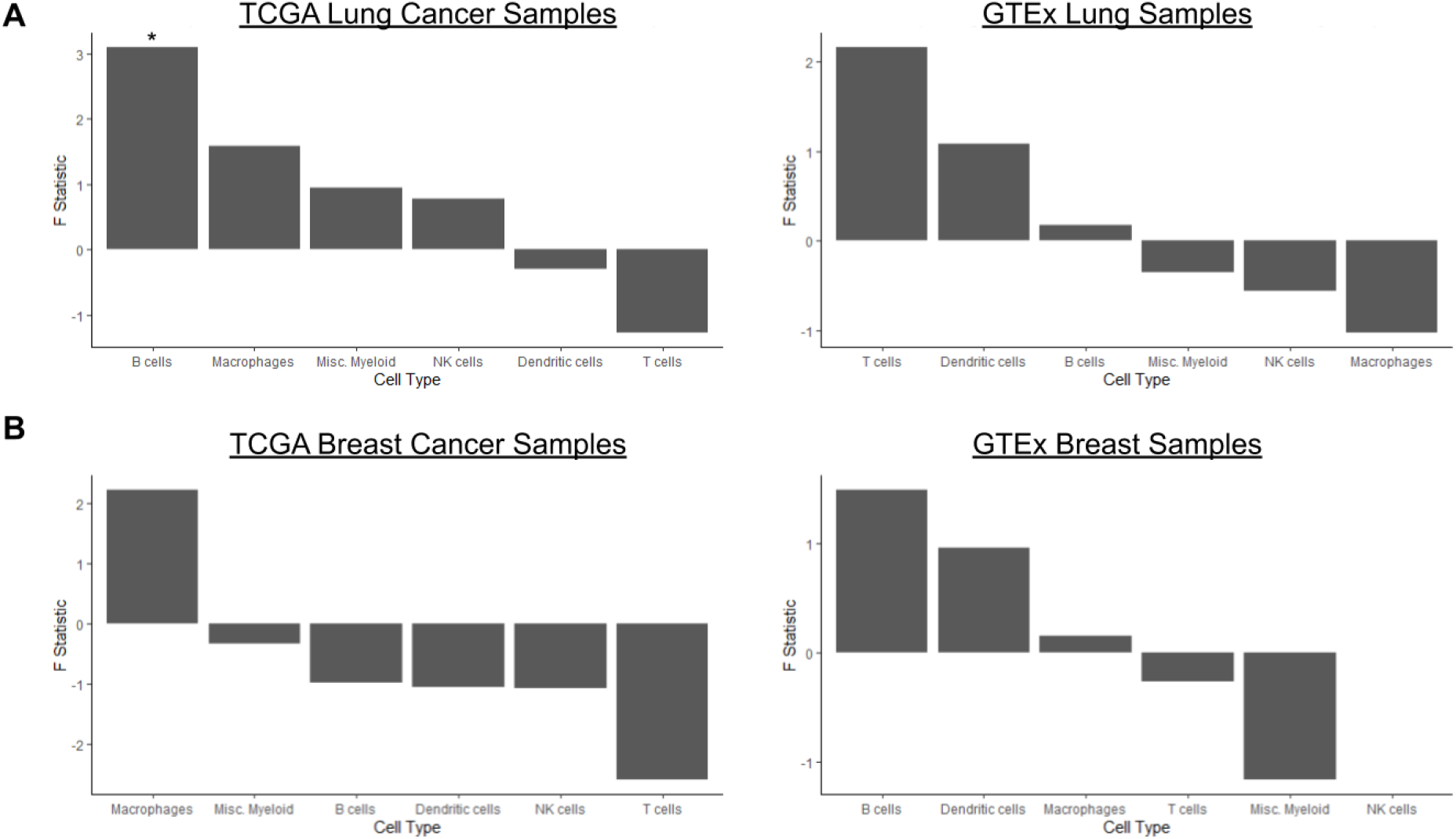
Association of immune cell type and age. The F statistic for the diagnosis age term of an ANOVA for each immune cell type (including age and sex as covariates) is used to compare **A** lung cancers to healthy lung tissue samples and **B** breast cancers to healthy breast tissue samples. * indicates a p-value less than 0.05.

**Supplemental Figure 5.**
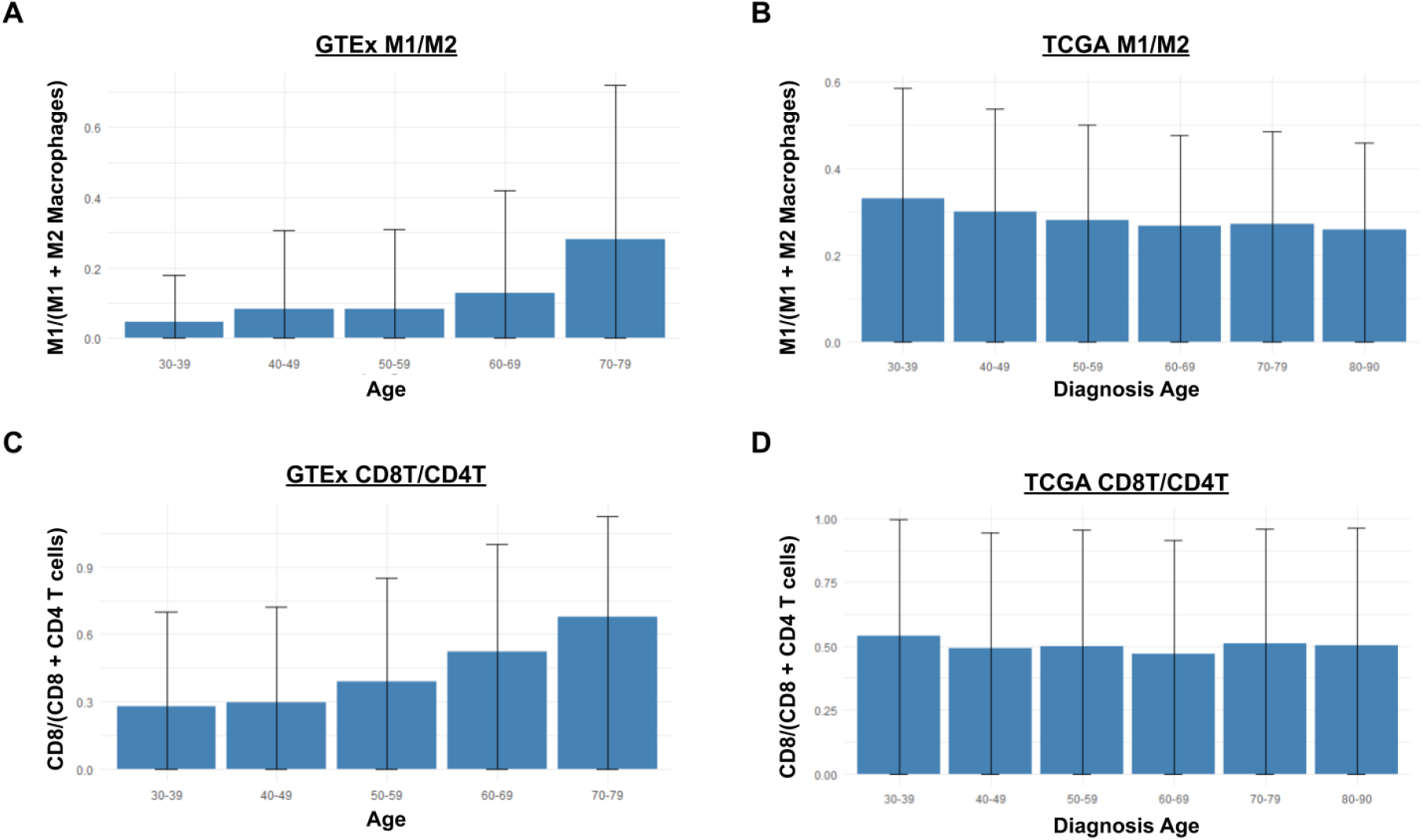
Comparing immune cell ratios of potential functional importance between TCGA and GTEx. **A-B** The proportion of M1 to M2 Macrophages in GTEx across tissues and TCGA pan-cancer, respectively. There is a significant increase in M1/M2 ratio in GTEx tissues with age, while there is no significant shift in TCGA. **C-D** Proportion of CD8 T cells to CD4 T cells with age in GTEx across tissues and TCGA pan-cancer, respectively. There is a significant increase in CD8/CD4 ratio in GTEx tissues with age, while there is no significant shift in TCGA.

**Supplemental Figure 6.**
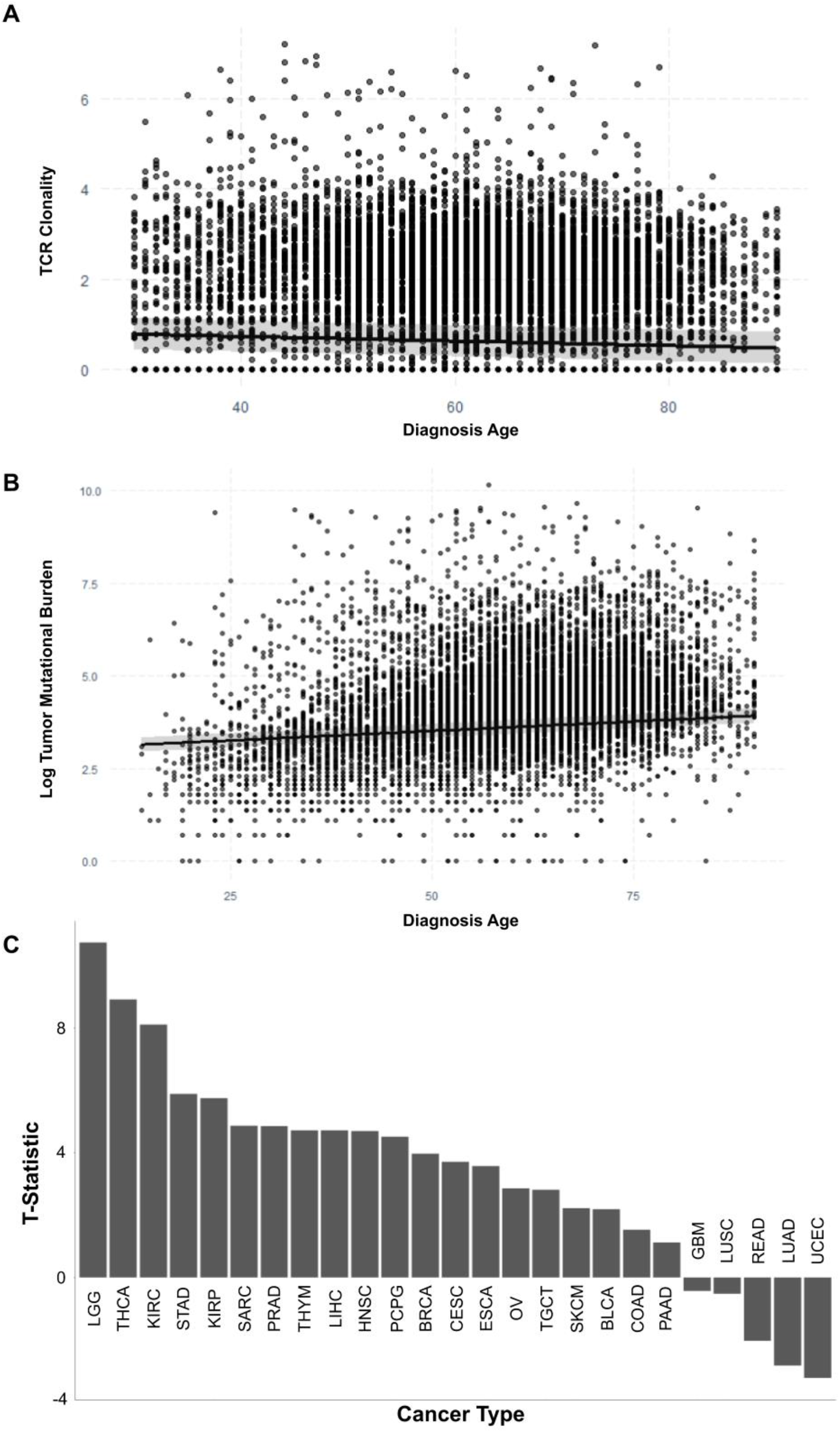
**A** Read normalized Shannon entropy of T cell receptor sequences plotted against age. The line fit to the data is a linear model of TCR clonality predicted by age, adjusted for sex and cancer type. **B** The log-normalized number of mutations found in each tumor sample is plotted against diagnosis age. The line fit to the data is a linear model of log-normalized number of mutations predicted by age, adjusted for sex and cancer type. **C** The T-statistic for the diagnosis age term of the linear model predicting the log normalized number of mutations for each cancer type.

**Supplemental Figure 7.**
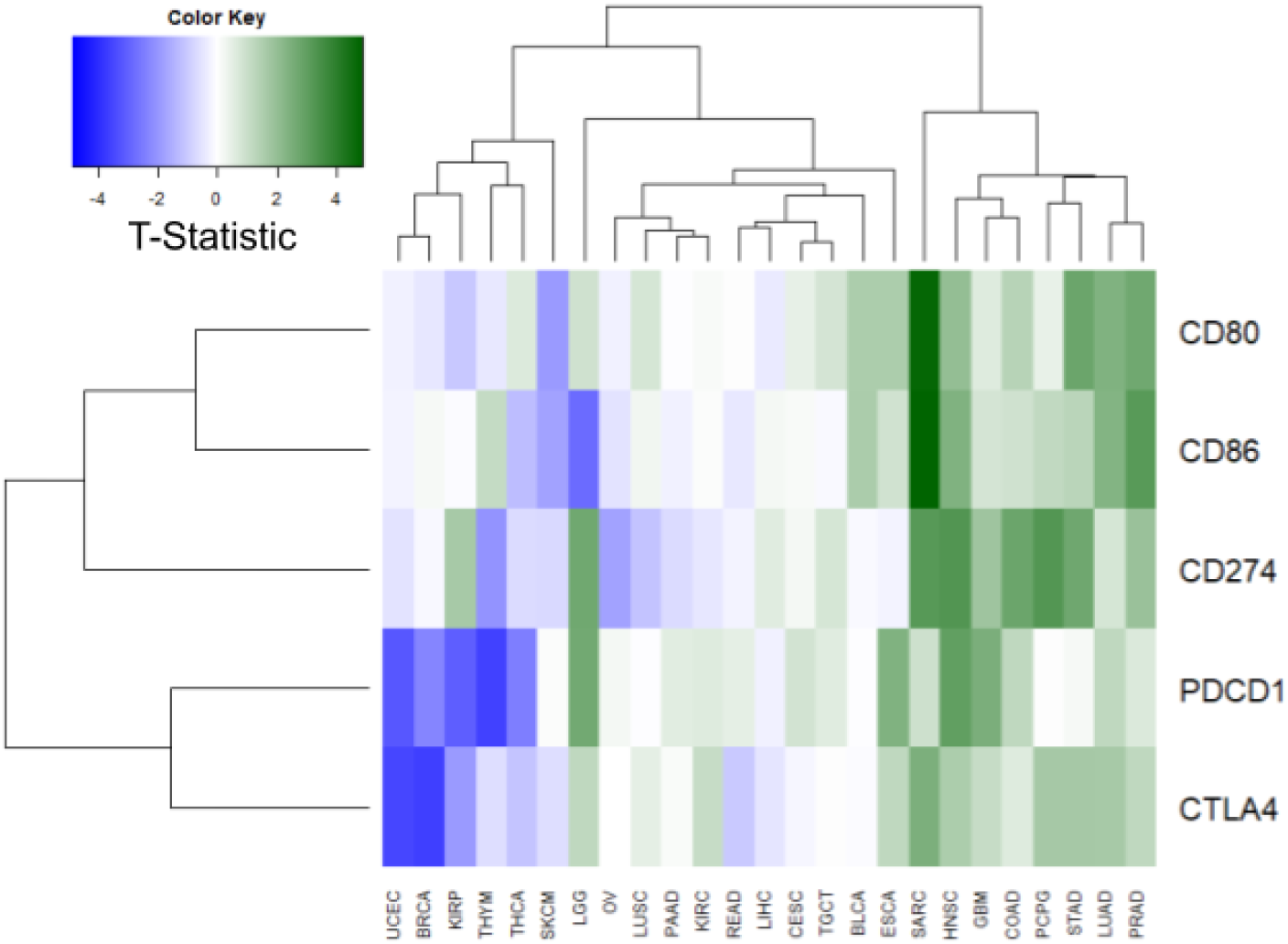
The association of immune checkpoint blockade genes with age across all cancer types with at least 100 samples in TCGA as determined by the t-statistic from limma-voom differential expression analysis.

**Supplemental Figure 8.**
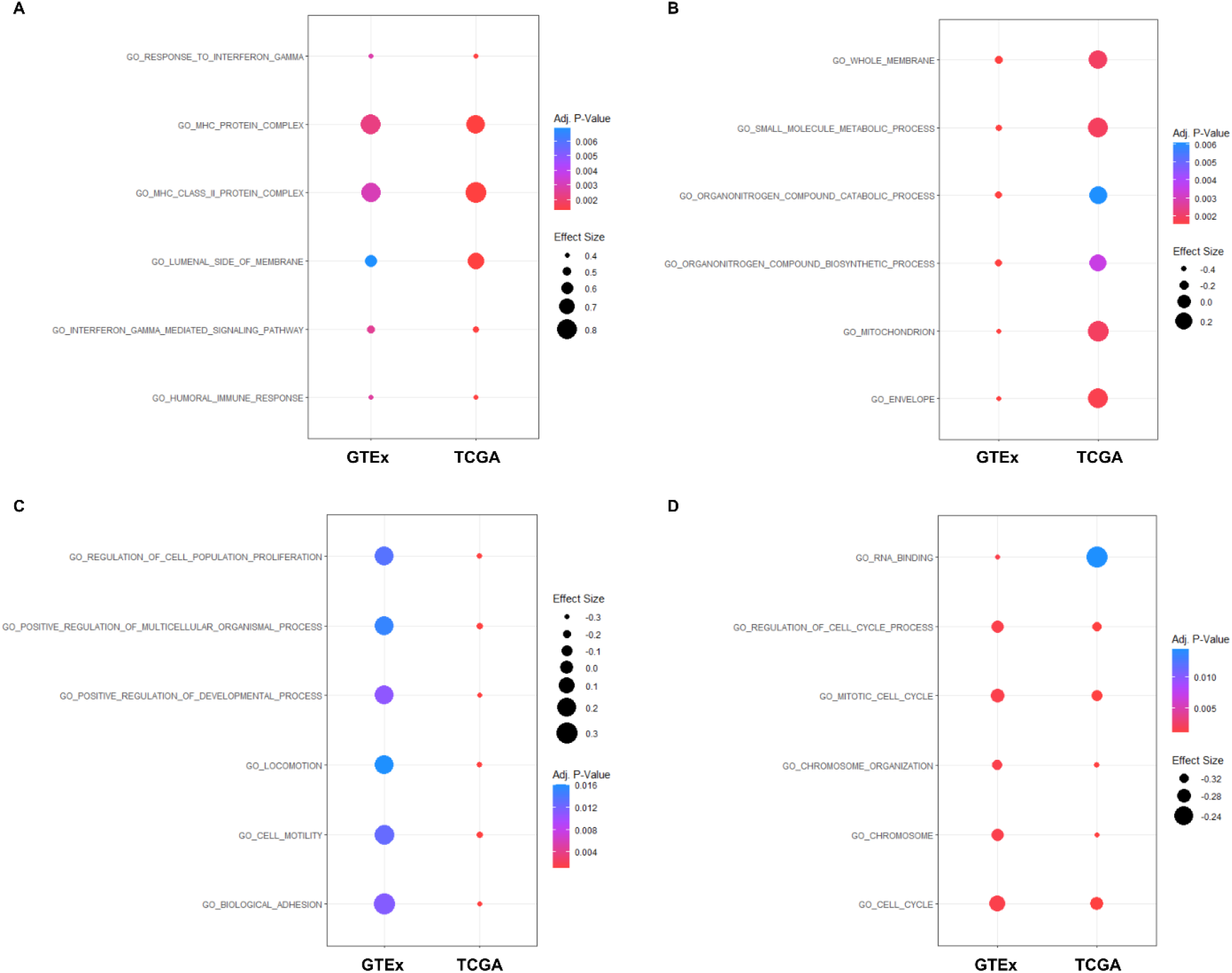
Dot plots representing genese set enrichment analysis for genes differentially expressed with age for both TCGA and GTEx. **A** Pathways significantly up in both TCGA and GTEx with age **B** Pathways significantly up in TCGA and down in GTEx with age **C** Pathways significantly up in GTEx and down in TCGA with age **D** Pathways significantly down in both TCGA and GTEx with age.

**Supplemental Figure 9.**
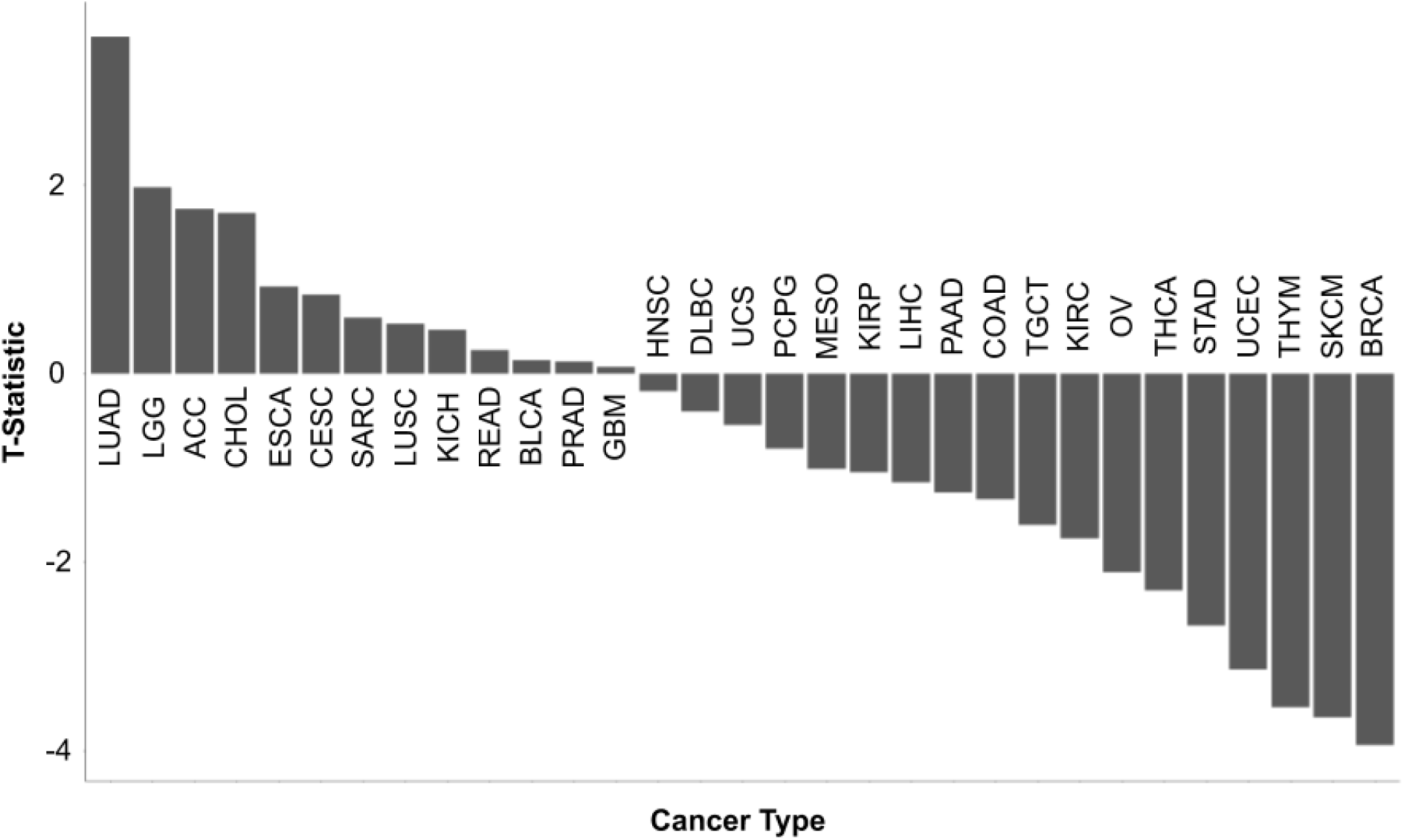
The association of TCR clonality and age is plotted for each cancer type in TCGA using the t-statistic from the diagnosis age term of a linear model fit to the TCR clonality of each cancer type, adjusting for sex as a covariate.

**Supplemental Figure 10.**
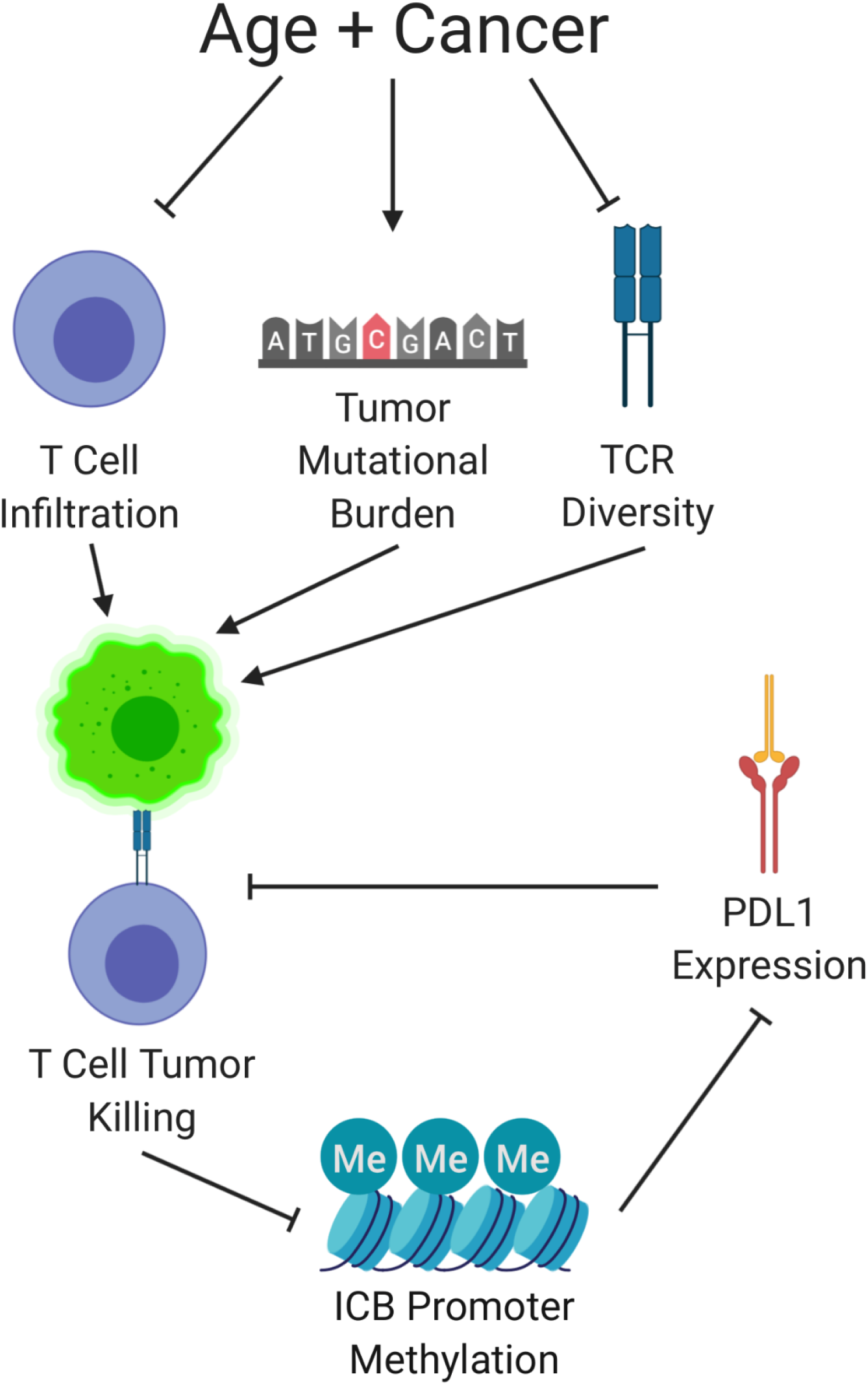
Hypothesized model for how the age-related effects we have observed affect T cell anti-tumor activity and expression of PDL1.

**Supplemental Table 1.** Immune cell types that were evaluated in relation to age for both TCGA tumor samples and GTEx normal samples.

| **Immune Cell Types** | **Immune Subtypes** |
| --- | --- |
| T cells | CD8 T cells, Naive CD4 T cells, Resting Memory Cd4 T cells, Activated Memory CD4 T cells, Follicular Helper T cells, Regulatory T cells, Gamma Delta T cells |
| Macrophages | M0 Macrophages, M1 Macrophages, M2 Macrophages |
| B cells | Naive B cells, Memory B cells, Plasma Cells |
| NK cells | Resting NK cells, Activated NK cells |
| Dendritic cells | Resting Dendritic cells, Activated Dendritic cells |
| Misc. Myeloid cells | Monocytes, Resting Mast cells, Activated Mast cells, Eosinophils, Neutrophils |

**Supplemental Table 2.** F-statistic, p, and q-values for each immune cell term in an ANOVA for the effect of age on the abundance of each immune cell type. Tissue and sex were included in the model as covariates.

|  | **F-statistic** | **p-value** | **q-value** |
| --- | --- | --- | --- |
| **T cells** | 0.887307 | 0.470706 | 0.564847 |
| **Macrophages** | 0.981287 | 0.416576 | 0.564847 |
| **B cells** | 5.835976 | 0.000117 | 0.00035 |
| **NK cells** | 19.34786 | 1.54E-15 | 9.27E-15 |
| **Dendritic cells** | 2.265888 | 0.060047 | 0.120095 |
| **Misc. Myeloid** | 0.647328 | 0.628807 | 0.628807 |

**Supplemental Table 3.** Number of genes differentially methylated and differentially expressed among immune related genes from TCGA samples. The genes with a canonical relationship between methylation and expression are in bold. The comparison is made both including and not including mutational burden as a covariate.

|  |  | Up Expressed | Down Expressed |
| --- | --- | --- | --- |
| Age, Cancer Type Adjusted | Up Methylated | 66 | **218** |
|  | Down Methylated | **113** | 94 |
| Mutational Burden Adjusted | Up Methylated | 62 | **135** |
|  | Down Methylated | **129** | 86 |

**Supplemental Table 4.** Basic demographic information used as covariates in the analyses conducted in this study comparing the TCGA and GTEx cohorts. *Age 20-29 was excluded from the TCGA patients collected (See Materials and Methods) and for comparison these individuals were excluded from the final analysis of GTEx data as well

|  | TCGA | GTEX |
| --- | --- | --- |
| Subjects | 8984 | 980 |
| Males | 4367 | 653 |
| Females | 4617 | 327 |
| Age 20-29* | N/A | 84 |
| Age 30-39 | 667 | 78 |
| Age 40-49 | 1226 | 153 |
| Age 50-59 | 2106 | 315 |
| Age 60-69 | 2453 | 317 |
| Age 70-79 | 1693 | 33 |
| Age 80-90 | 504 | 0 |

**Supplemental Table 5.** Number of RNA-seq samples from each tissue in GTEx data used.

| **Tissue** | **Number**<br>**of**<br>**Samples** |
| --- | --- |
| Adipose<br>Tissue | 1204 |
| Adrenal<br>Gland | 258 |
| Bladder | 21 |
| Blood | 929 |
| Blood<br>Vessel | 1335 |
| Brain | 2642 |
| Breast | 459 |
| Cervix<br>Uteri | 19 |
| Colon | 779 |
| Esophagus | 1445 |
| Fallopian Tube | 9 |
| Heart | 861 |
| Kidney | 89 |
| Liver | 226 |
| Lung | 578 |
| Muscle | 803 |
| Nerve | 619 |
| Ovary | 180 |
| Pancreas | 328 |
| Pituitary | 283 |
| Prostate | 245 |
| Salivary Gland | 162 |
| Skin | 1809 |
| Small Intestine | 187 |
| Spleen | 241 |
| Stomach | 359 |
| Testis | 361 |
| Thyroid | 653 |
| Uterus | 142 |
| Vagina | 156 |

**Supplemental Table 6.** Number of RNA-seq samples from each TCGA study in the TCGA data used in this work.

| **TCGA**<br>**Cancer**<br>**Type**<br>**Acronym** | **Number**<br>**of**<br>**Samples** |
| --- | --- |
| ACC | 64 |
| BLCA | 405 |
| BRCA | 1056 |
| CESC | 270 |
| CHOL | 35 |
| COAD | 272 |
| DLBC | 45 |
| ESCA | 180 |
| GBM | 148 |
| HNSC | 507 |
| KICH | 61 |
| KIRC | 508 |
| KIRP | 276 |
| LAML | 158 |
| LGG | 433 |
| LIHC | 347 |
| LUAD | 479 |
| LUSC | 474 |
| MESO | 86 |
| OV | 292 |
| PAAD | 177 |
| PCPG | 157 |
| PRAD | 480 |
| READ | 88 |
| SARC | 244 |
| SKCM | 411 |
| STAD | 403 |
| TGCT | 74 |
| THCA | 433 |
| THYM | 116 |
| UCEC | 169 |
| UCS | 57 |
| UVM | 79 |

